# Hot cue: Physiologically controlled release from an in situ forming liposomal depot

**DOI:** 10.1101/2025.10.22.683980

**Authors:** Remo Eugster, Simone Aleandri, Julia Bassila, Davide Bochicchio, Laura Baraldi, Belinda Hämmerle, Stefan Schürch, Martina Vermathen, Peter Vermathen, Giulia Rossi, Alessandra Bergadano, Paola Luciani

## Abstract

Poor treatment adherence, often referred to as the “silent epidemic”, is a growing global issue that significantly contributes to preventable illness, premature death, and rising healthcare costs. Among compliance-enhancing strategies, controlled-release depots designed for intermediate treatment cycles represent a suitable approach, particularly in contexts where frequent dosing is impractical. Current long-acting injectable formulations, though, are often hindered by challenges in injectability, delayed onset, and complex manufacturing. Here, we present a thermoresponsive, in situ forming liposomal depot (TILD) designed to modulate the release of membrane-associated drugs following subcutaneous administration. Using buprenorphine as a model analgesic, we demonstrate that TILD responds to subcutaneous divalent cations with immediate surface-drug release and to body temperature with sustained diffusion through a fluidized bilayer. Molecular simulations guided the system design, and structural and colloidal characterizations validated its responsiveness to physiological cues. In vivo, TILD formed stable depots and maintained therapeutic drug levels for up to four days in both rats and Beagle dogs. Pharmacodynamics studies in rats confirmed the delivery of prolonged analgesia.

## Introduction

Lack of treatment adherence, often described as a ‘silent epidemic’, continues to undermine outcomes across human and veterinary medicine.[1] Frequent drug administration remains a major challenge in clinical care, particularly in populations with limited compliance, such as geriatric, incapacitated, or veterinary patients.[2–5] In human and veterinary medicine alike, complex treatment regimens often require multiple daily drug administrations, as seen in the management of post-surgical pain.[3,6–8] These regimens cause distress and logistical challenges in patients, caregivers and animals, leading to poor compliance and suboptimal patient outcomes.[3,6] In this matter, the subcutaneous (s.c.) route has gained prominence due to its ease of administration, potential for self-injection, and suitability for sustained drug release for a broad spectrum of therapeutics, from small molecules to complex biotherapeutics.[9,10]

Extended-release drug formulations could substantially reduce the burden of frequent administration, but their clinical translation is frequently hindered by complex manufacturing, poor scalability, and difficult administration stemming from physical or handling limitations of the formulation.[3,11] Parenteral depots often require large volumes, high viscosity, and aseptic processing - factors that limit their practicality in real-world settings.[3,12] These challenges underscore the need for new delivery systems that offer extended pharmacokinetics without compromising usability or manufacturability.[1]

Liposomes are well-established, versatile drug carriers capable of enhancing stability, bioavailability, and pharmacokinetics of a wide variety of therapeutic agents (Fig 1A).[13] With tailored lipid compositions, liposomes can encapsulate a broad range of drug molecules and enable controlled release.[13] However, most long-acting systems fail to provide both rapid onset and sustained therapeutic levels, often requiring an initial bolus dose or relying on complex external triggers such as heat, light, or hardware-assisted activation.[3,11] This limits their practicality and consistency, particularly in outpatient or veterinary settings.[14,15] A drug delivery system that forms a depot in situ after injection, remains localized at the administration site, and releases its payload in a controlled, dual-mechanistic manner - combining immediate and prolonged effects - could address critical barriers in clinical and translational medicine.[2,16]

**Fig 1.**
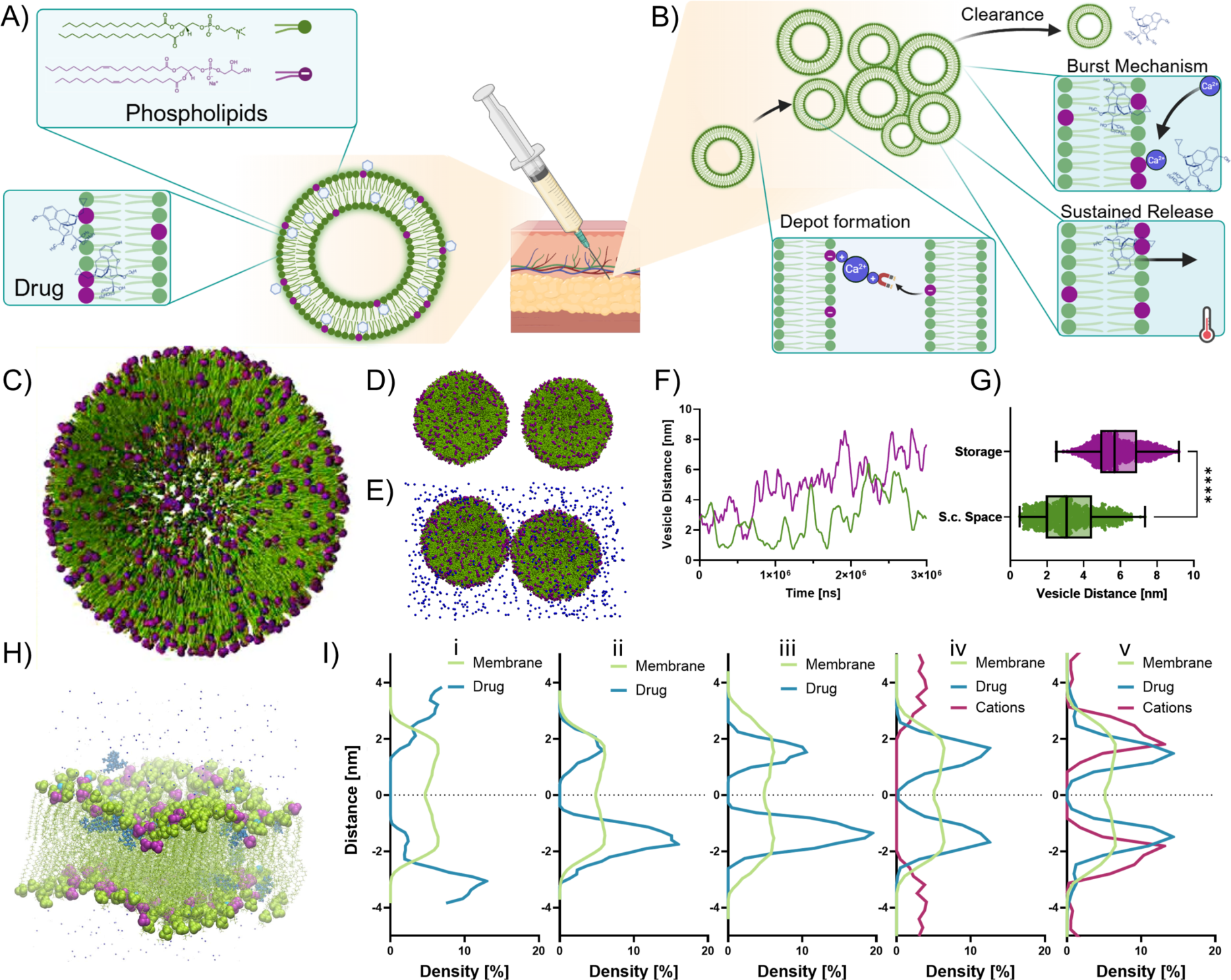
Design and trigger-responsive concept of the thermoresponsive in situ forming liposomal depot (TILD) system. (A) Liposomal composition and drug encapsulation (B) Depot formation and drug release concept induced by physiological triggers. Molecular dynamics simulations of conceptual liposomal drug delivery (C) Coarse grained simulation of liposomal vesicle (D) Snapshot of two liposomes of the TILD system in simulated storage conditions with a notable distance between the vesicles. (E) Snapshot of said liposomes within a simulated subcutaneous (s.c.) space. Demonstration of divalent cations facilitating bridges between the two vesicles, pulling them closer together and aggregating while maintaining their vesicle shape. (F) Quantification of the distances between the vesicles at different simulation conditions, demonstrating aggregation driven by divalent cations. (G) Scatter plot representing average distance between vesicles over simulation time (after 1 µs of equilibration). (H) Simulation snapshot of liposome membrane, with all-atom resolution, within an approximated s.c. environment in the presence of divalent cations illustrating the position of drug molecules (blue) within the membrane composed of HSPC lipids (green), DOPG lipids (green tails, purple heads), and physiological NaCl. (I) Density along z-axis of membrane, drug molecules and physiological cations during consecutive simulations illustrating position and movement of molecule groups within the simulation box. i. start of drug loading (65 °C); ii. Drug incorporated into membrane iii. Cool down to storage temperature (4 °C); iv. Warm up to body temperature and exposure to simulated s.c. space; v. After exposition and surface associated drug release.

Here, we introduce a **thermo**responsive **i**n situ forming **l**iposomal **d**epot (TILD) for the treatment of post-surgical pain that combines ease of administration with precise spatiotemporal drug release. Upon subcutaneous injection, TILD transitions from a nanoparticle dispersion to a depot structure triggered by physiologically present ions (Fig 1B). This enables an initial burst release phase triggered by divalent cation-mediated displacement of membrane-associated drug molecules, followed by a second phase of sustained release driven by temperature-induced bilayer permeability.

Through molecular dynamics simulations and experimental validation, we demonstrate that drug-lipid interactions are controllable by temperature and ionic conditions. Cations present in the s.c. space displace drug molecules from the vesicle surface while increasing temperature induces structural changes within the bilayer, enabling sustained diffusion. In vivo, TILD provides therapeutic plasma concentrations within minutes of administration and maintains efficacy over multiple days in rats and with a successful pharmacokinetic and tolerability evaluation in beagle dogs. The modular design and mechanistic understanding of the system provide a basis for potentially adapting this approach to a range of physicochemically similar drug molecules and therapeutic contexts. While drug-specific adjustments to the system might be needed, computational tools can provide a fast, generalizable framework for their design.

## Results & Discussion

### Mechanistic Computational Design

The design of TILD was guided by three key principles to achieve a dual-mechanism release upon subcutaneous administration. First, we leveraged the net surface charge of the vesicles to promote in situ depot formation through ion-bridging interactions with physiological polyvalent cations, ensuring retention at the injection site.[17,18] Second, we sought to enable a rapid release phase by associating a fraction of a basic, positively charged drug to the negatively charged 1,2-dioleoyl-*sn*-glycero-3-phosphoglycerol (DOPG) headgroups at the liposome surface.[19] Upon injection, divalent cations present in the s.c. space are expected to displace these surface-bound drug molecules by competing for the same anionic lipid headgroups. Third, we engineered the lipid composition, selecting hydrogenated soy phosphatidylcholine (HSPC) in combination with DOPG, to undergo a phase transition at physiological temperature, enabling a sustained release of the remaining drug fraction, embedded within the bilayer, due to its amphiphilic nature.[19] Using a computational approach, we predicted and fine-tuned these design parameters with an understanding at atomistic level, reducing experimental trial-and-error and accelerating formulation development.[20–23]

To understand and validate these mechanisms, we performed atomistic and coarse-grained molecular dynamics simulations (Fig 1).[20,21] These simulations allowed us to resolve the localization, orientation, and mobility of drug molecules within the liposomal bilayer and enabled us to assess the influence of s.c. ionic conditions on vesicle behaviour and aggregation dynamics.[23] For these studies, we used buprenorphine, a clinically relevant opioid commonly used for the treatment of post-surgical pain,[24–27] chosen as a model compound to evaluate both immediate and sustained analgesic delivery through the TILD system, aiming at closing a therapeutic gap in both, veterinary and human medicine.[26–29]

Coarse-grained simulations provided insight into depot formation (Fig 1C).[30] Under simulated storage conditions, liposomes remained well dispersed (Fig 1D). To approximate the s.c. environment, Ca^2+^ was included at physiological concentrations during model setup (Fig 1E).[31,32] Upon exposure to physiological ion concentrations, vesicles moved closer and formed stable aggregates via ion bridging, without membrane fusion or loss of vesicle integrity (Figs 1F-G). This aggregation behaviour mimics the depot formation observed in vivo, in which vesicles remain localized, allowing for sustained drug diffusion from the depot site.[18] Atomistic simulations of the drug-loaded membrane were executed over multiple consecutive simulations resembling its path from production to application. First, the assembly and drug loading of the membrane resembling liposome production was simulated (Fig 1H). During simulated production, it was revealed that the drug buprenorphine preferentially interacts with the negatively charged DOPG headgroups within the bilayer, orienting so that ionizable moieties remain near the membrane surface (Fig 1I-i & ii and Supporting Fig S1). In a second stage, the vesicle membrane was cooled to storage temperature. Cooling the formulation to storage temperature causes buprenorphine molecules to embed deeper into the hydrophobic core at a consistent orientation, becoming shielded and immobilized (Fig 1I-iii and Supporting Fig S1).[32] In a last step, the drug-loaded membrane was warmed to body temperature and exposed to an approximated simulation of the s.c. space in terms of divalent cations (Fig I-iv).[32,33] When simulating the s.c. environment, we observed that divalent cations, particularly Ca²⁺, displaced surface-associated drug molecules by competitively binding to DOPG headgroups, resulting in an envisioned burst release (Fig 1I-v, Supporting Fig S1). Additionally, exposure to physiological temperatures promotes bilayer fluidization, restoring molecular mobility and facilitating sustained release.[34]

Together, these simulations elucidate how TILD leverages lipid composition and environmental responsiveness to enable both rapid onset and extended drug release. The mechanism should facilitate a dual-mechanism release, highlighting the critical role of DOPG- cation interactions in modulating immediate drug availability. Importantly, the computational approach provides a flexible development platform that could be rapidly adapted to study other small-molecule drug candidates with similar physicochemical properties and lipid compositions, streamlining the development of tailored delivery on this potential platform.

### Physicochemical Validation

While molecular simulations provided crucial insight into the structural and mechanistic basis of TILD and allowed a streamlined formulation design, experimental validation is essential to confirm that these functions persist under real-world conditions, as with all computational approaches.[22] Critical features predicted computationally were assessed experimentally to validate both the design rationale and the functional responsiveness of TILD under physiological conditions.

Localization of buprenorphine within the bilayer and at the liposomal surface, as predicted by simulations, was confirmed experimentally by small-angle X-ray scattering (SAXS) and nuclear magnetic resonance (NMR).[35,36] SAXS revealed increased electron density at both headgroup and bilayer core regions, indicating that buprenorphine partitions into both domains (Fig 2A). At elevated temperatures (38 °C), electron density near the headgroups increased, suggesting drug drift toward the membrane surface. Upon exposure to physiological levels of divalent cations, this density decreased - consistent with cation-driven displacement of surface-associated drug - and was accompanied by structural changes in lipid packing, supporting the mechanism observed in atomistic simulations (Fig. 1I).[36] Complementary NMR analyses indicated interactions with both headgroup and tail moieties, with partial drug dissociation upon cation exposure, supporting cation-mediated release of surface-associated drug in an initial release phase (Supplementary Fig S2).[37] These findings align with simulation data, showing surface interactions driven by electrostatic attraction of the drug to negatively charged DOPG headgroups. The favourable interaction of buprenorphine with both the liposomal surface and the lipid bilayer interior resulted in an encapsulation efficiency of ∼95% at a drug-to-lipid ratio of 1:20 (Fig 2B). Although slightly lower ratios may improve encapsulation marginally, they require significantly more lipid to achieve therapeutic doses, complicating formulation and administration due to higher viscosities.[13] Importantly, the distribution of buprenorphine across both surface-bound and bilayer-embedded states supports efficient encapsulation and enables distinct release mechanisms in response to physiological conditions.

**Fig 2.**
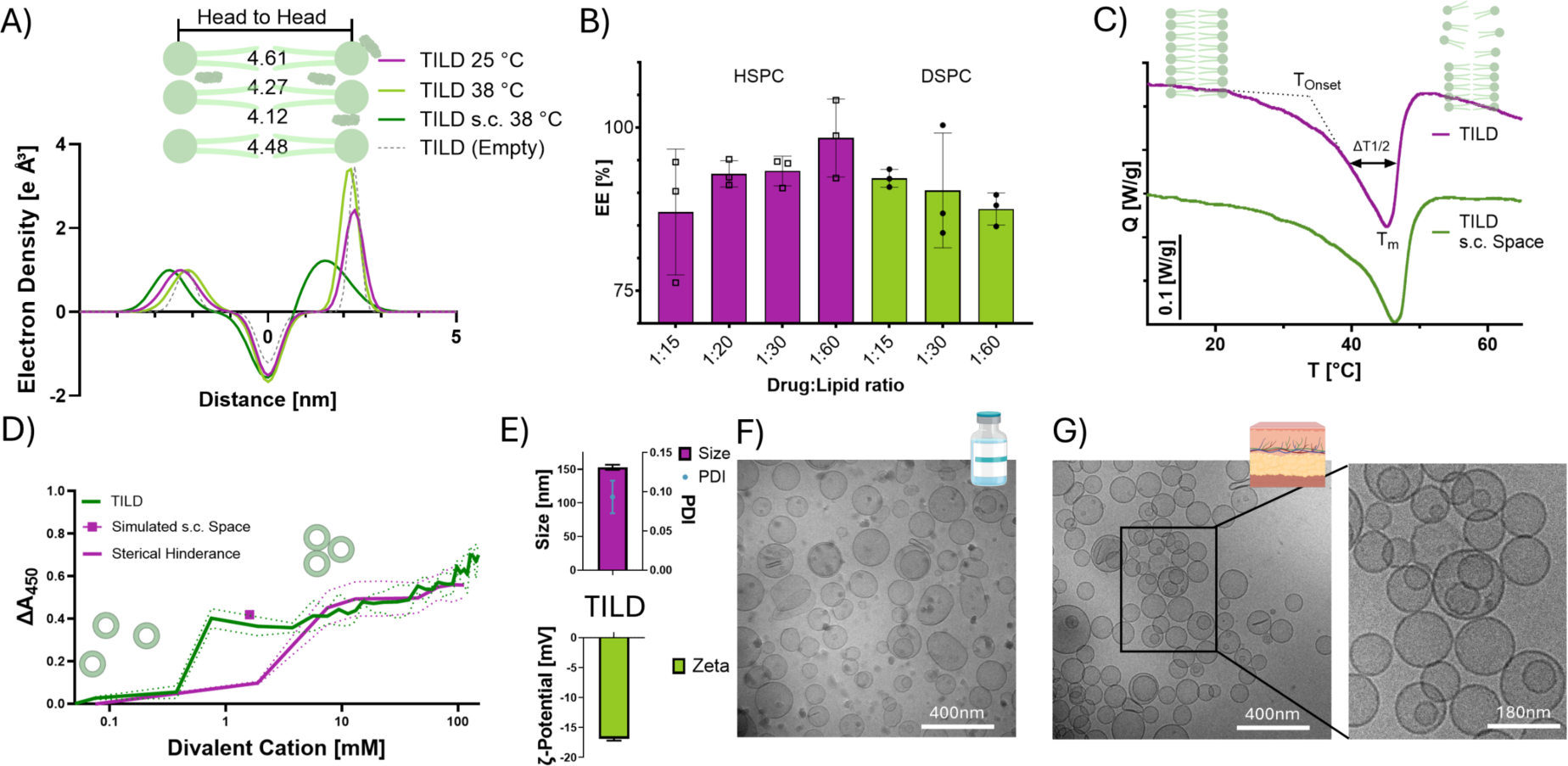
Physicochemical characterization of the drug delivery system TILD (A) Electron density profiles obtained from SAXS measurement at 25 °C and 38 °C and with divalent cations at physiological levels. (B) Encapsulation efficiency (EE%) of buprenorphine within TILD and liposomes where HSPC has been replaced with DSPC at different drug to lipid ratios. (n=3; mean±SD). (C) Thermograms of vesicles indicating phase transition temperature T_m_ and its onset. (D) Depot formation profiles of TILD along an increasing concentration gradient of divalent cations. A mimicked s.c. space marks a point of depot formation at physiological state (∼1.6 mM Ca²⁺). PEGylated liposomes, which facilitate sterically hindrance, were used as control. (n=3; mean±SD). (E) Vesicle characterization in terms of hydrodynamic diameter, PDI, and surface charge (zeta potential). (n=3; mean±SD) (F) Cryo-TEM images of vesicles under mock storage conditions (G) and vesicles exposed to physiological concentrations of divalent cations (∼1.6 mM Ca²⁺), mimicking the subcutaneous environment, exhibit surface adhesion, forming grape-like aggregates while preserving their membrane integrity.

Computational predictions further indicated that the embedded drug population becomes more deeply embedded in the membrane at storage temperatures, thereby enhancing stability (Fig 1I; Supporting Fig S1). In contrast, physiological temperatures induce structural changes that facilitate drug diffusion and enable sustained release.[38] Differential scanning calorimetry (DSC) revealed the membrane’s thermal responsiveness, with the phase transition occurring near body temperature (34−38 °C). This transition, essential for enabling drug diffusion and release, was highly dependent on the inclusion of DOPG and its specific combination with HSPC (Supporting Table S1).[38] Analysis of the cooperativity of the bilayer confirmed that both the incorporated drug and divalent cations at physiological concentrations reduced the sharpness of this transition. The temperature width at half-height (ΔT_1/2_) increased from 7.37 K for the empty nanovesicles to 8.9 K with the drug, and further to 9 K with cations. This reduced cooperativity, indicated by a broader phase transition peak, is a favourable design feature. It shifts the drug release profile from a sharp, burst-like event to a more gradual, sustained process, allowing for the precise tuning of the temperature-driven release kinetics. To further validate the role of vesicle-level behaviour in depot formation, we examined how the colloidal properties of TILD respond to physiologically relevant cues. Dynamic light scattering (DLS) and turbidity measurements demonstrated that TILD vesicles form stable, micron-scale aggregates upon exposure to divalent cations, such as calcium, which are abundant in the s.c. space (Fig 2D and Supporting Fig S3). The replacement of DOPG with other lipids was not viable, as DOPG is critical for formulation stability, surface drug coupling, and depot formation through ionic interactions with divalent cations (Supporting Fig S3). These findings are consistent with the simulated ion-bridging mechanism and support the concept of depot formation via cation-mediated vesicle clustering.

Cryo-electron microscopy (Cryo-TEM) provided direct visual confirmation of this feature. Under storage conditions, vesicles remained dispersed and well-defined (Fig 2F). Upon exposure to physiological calcium concentrations (∼1-2 mM),[31,32] vesicles clustered into dense but intact networks, mimicking the depot morphology (Fig 2G). This stable network ensures localized retention and is critical for minimizing systemic clearance and extending the duration of drug availability.[18]

Together, our experimental observations confirm that TILD performs as designed: divalent cations first displace the surface-associated drug through electrostatic competition, then drive vesicle aggregation and depot formation, while subsequent temperature elevation increases bilayer fluidity to enable sustained release of membrane-embedded drug - all while maintaining formulation stability under storage conditions (Supporting Fig S4). The alignment between computational predictions and empirical validation reinforces the robustness of the design and its translational potential. Beyond mechanistic design and physicochemical validation, formulation scalability was addressed using a microfluidic manufacturing strategy, supporting the feasibility of translating TILD to relevant production platforms (Supporting Fig S5).[22,39]

### In vivo translation

Having established that TILD responds predictably to ionic triggers and temperature, we next assessed its functional performance in vitro and in vivo to confirm that the dual-mechanism release function translates into therapeutic outcome.

To experimentally confirm the release behaviour described earlier, in vitro drug release studies using horizontal diffusion cells were conducted under conditions mimicking the s.c. environment, demonstrating a release profile driven by two mechanisms (Fig 3A). Upon exposure to physiological conditions, an initial burst of drug release was observed, corresponding to the displacement of surface-associated drug by divalent cations (Fig 3A), resulting in a similar drug release profile as the free drug for the first ∼10%. This was followed by a prolonged release phase, driven by temperature-induced membrane fluidization and drug diffusion from within the bilayer. This transition became apparent as the release rate diverged from that of the free drug control, reflecting the onset of temperature-induced diffusion through the lipid bilayer. The biphasic model (R² = 0.997) provided a good fit, though it offered only a marginal improvement over the monophasic model when accounting for the added complexity, as indicated by comparisons using the Akaike and Bayesian Information Criteria. Nevertheless, control formulations lacking responsive elements - such as neutral vesicles or pre-aggregated vesicles - exhibited either premature release or poor diffusion, underscoring the role of lipid composition and depot formation in achieving the desired release kinetics (Supporting Fig S6).

**Fig 3.**
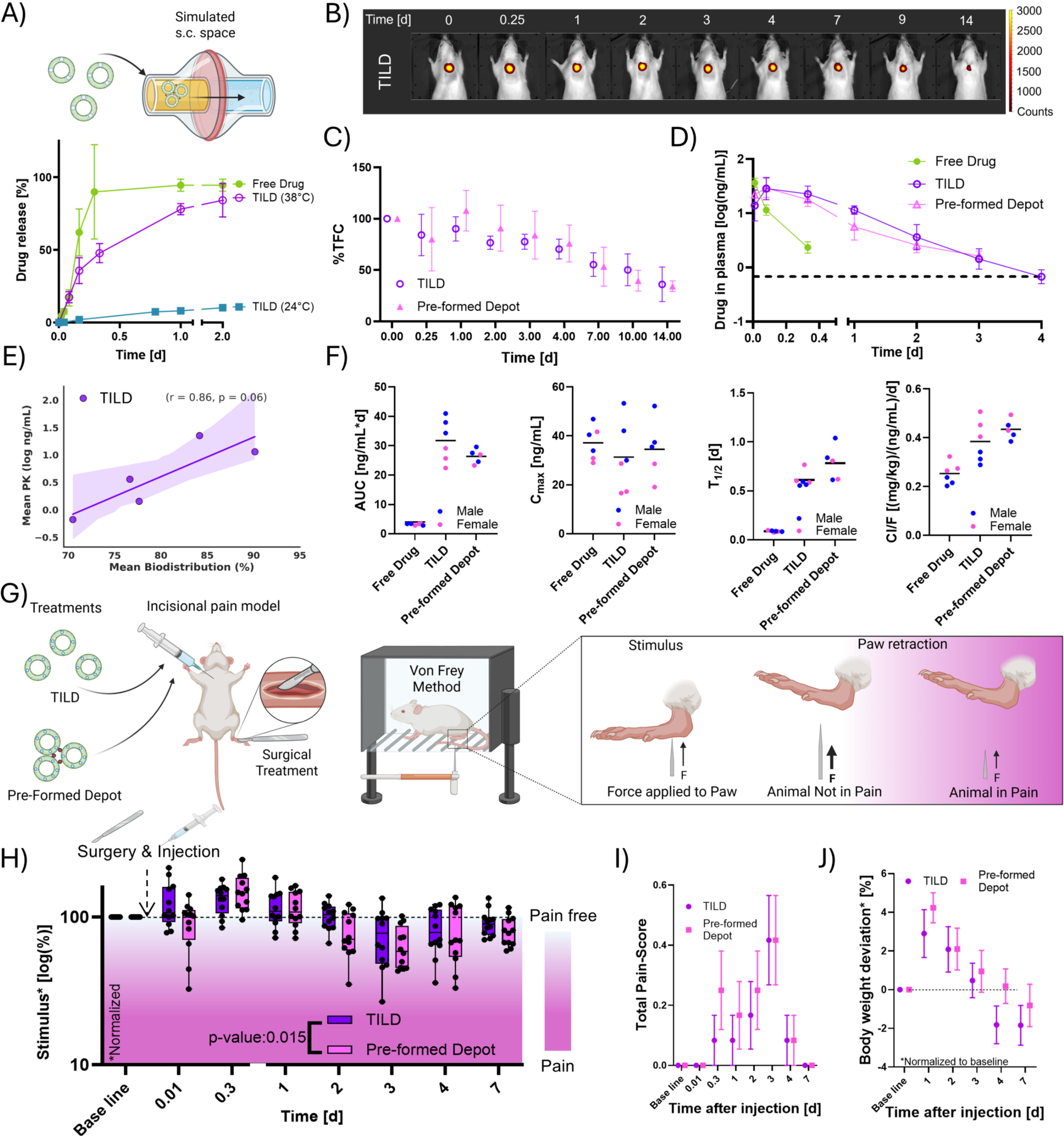
Functional validation of TILD (A) Drug in vitro release comparing free drug against TILD at different temperatures, and divalent cations to simulate the s.c. space and storage conditions. (B) Representative images of TILD forming a depot structure in s.c. space after injection (C) Quantification of total fluorescence counts (TFC) normalized to T_0_ at injection vs time of TILD and pre-formed depot (n=4; mean ±SD). (D) Pharmacokinetics of free drug at clinical dose as well as TILD and pre-formed depot compensating for 12 clinical doses of the fast-acting drug with the dashed line indicating lower bound of therapeutic window (n=6; mean ±SD). (E) Correlation of Biodistribution vs. PK for TILD with Pearson correlation coefficient (F) Pharmacokinetic parameters by sex (n=6; mean). (G) Pain assessment in a postoperative incision model using a von Frey method. (H) Post-surgical pain stimulus assessed via von Frey measurements for TILD and pre-formed depot (n=12; boxplot incl. mean and quartiles) (Stat.: Linear mixed-effects model). (I) Pain score indicating animal distress for each tested formulation (See Supporting Information for details on pain score) (n=12; mean ±SEM). (J) Body weight of animals treated with different formulations (n=12; mean ±SEM).

These findings were mirrored by comprehensive in vivo studies. Imaging studies visually confirmed the formation of the fluorescently labelled system in vivo (Fig 3B). TILD remained localized at the injection site for several days, closely mirroring the behaviour of pre-formed depots, thereby reducing systemic clearance and ensuring sustained drug availability (Fig 3C).[18] Depot integrity began to be compromised after day four, aligning well with the observed pharmacokinetic profile and minimizing the risk of long-term tissue accumulation. Unlike conventional sustained-release systems that are preformed, often viscous, and rely solely on passive diffusion or polymer degradation,[3] TILD forms a responsive depot at the injection site through well-defined physicochemical interactions.

Pharmacokinetic studies confirmed that TILD rapidly achieved therapeutic plasma concentrations following subcutaneous injection - matching the early onset of the free drug - while also exhibiting depot-mediated sustained release afterwards (Fig 3D). TILD maintained plasma drug levels within the therapeutic window (0.75 ng/mL to >5 ng/mL) for up to 3–4 days, in contrast to the short half-life of the conventional free drug formulation.[40] Notably, a single dose of TILD has the capability to replace the need for approximately twelve consecutive injections of free drug in the rat model, underscoring its potential to reduce treatment burden and improve adherence significantly.[3,6,7,41] Furthermore, the persistence of the drug appears to be driven by the depot system, as indicated by a Pearson correlation between the clearance phases of the depot-associated nanoparticles and the drug in plasma (Fig. 3E). This extended exposure reflects the intended depot function and aligns with the computational and in vitro release profiles. Notably, TILD achieved this without peak plasma concentrations (C_max_) exceeding those of the free drug, suggesting a reduced risk of systemic adverse effects (Fig 3F).[42] Also, the area under the curve (AUC) clearly indicates TILD’s capability to deliver its payload and compensate for 12 consecutive injections of the free drug and clear it later in a retained fashion (Fig 3F).

In pharmacodynamic studies using a rat model of post-surgical pain,[43] TILD demonstrated fast-acting and sustained analgesia following a single dose (Fig 3G). For ethical reasons, a no-analgesia group was not included, as withholding pain treatment following surgery is known to cause significant distress and hypersensitivity.[44,45] However, extensive prior studies have shown that untreated animals exhibit markedly reduced withdrawal thresholds, consistent with severe postoperative pain.[43,46] Similarly, a free drug control was not included, as its analgesic effect in this model is well established to last only 6–8 h.[43,46] Besides that, compared to pre-formed depots, TILD provided faster onset and extended efficacy for up to 72 h - addressing a key challenge in postoperative pain management (Fig 3H).[8] Also, just minor animal scoring (Supporting Information) changes and deviation of bodyweight from study start were observed, indicators for post-operative pain, further demonstrating TILDs efficacy (Fig 3I-J).[47] In addition, TILD requires no pre-aggregation steps and maintains a lower viscosity compared to pre-formed depots. This improves injectability by enabling faster, simpler administration through standard subcutaneous needles.[48] The drug effect was designed to subside around day three, a window that aligns with routine rechecks, allowing for clinical reassessment without masking potential symptoms.[29,49]

### Clinical translation for veterinary use

The translational potential and tolerability of the system were evaluated in beagle dogs, a large-animal model with high anatomical and metabolic similarities to humans, serving as a preclinical bridge toward future clinical studies.[50] Following subcutaneous administration, TILD exhibited a rapid onset within minutes and sustained plasma concentrations of buprenorphine over 48–72 h (Fig. 4A). In contrast, extrapolated pharmacokinetic data from Nunamaker et al. for the current clinical standard - a single subcutaneous dose of non-controlled buprenorphine at 0.02 mg/kg - shows therapeutic levels persisting for only 4-6 hours.[51] Compared to the rat model (Fig. 3), drug plasma levels in beagle dogs remained at a more consistent plateau for approximately two days before declining, suggesting that the larger subcutaneous compartment may enhance depot stability. The AUC averaged ∼6 ng·d/mL, supporting prolonged systemic exposure (Fig 4B). Notably, the C_max_ of TILD (∼3 ng/mL) was substantially lower than that reported for the fast-acting formulation (∼19.6 ng/mL), potentially reducing the risk of concentration-related side effects (Fig 4B).[51] Although the pharmacokinetics were evaluated in a small sample size that cannot be generalized to the broader dog population, clearance was consistent across subjects, indicating favourable release kinetics (Fig 4B). The mean residence time (MRT), which reflects how long the drug remains in systemic circulation, exceeded 36 h - consistent with sustained release and indicative of extended analgesic coverage (Fig. 4B). The formulation was well tolerated, with no unexpected adverse events (Supporting Information). Mild, transient effects - light sedation, reduced heart rate, and lower body temperature - were observed, aligning with the known pharmacodynamic profile of buprenorphine (Fig. 4C).[51] This additional layer of validation of the release kinetics underscores the readiness of TILD for further development in both human and veterinary applications. This is particularly important considering therapeutic opportunities on the rise in the companion animal landscape.[52] Large-animal data is hence crucial at this stage, providing translational confidence in anatomical compatibility, injectability, and pharmacological behaviour in species closer to humans.[53–55]

**Fig 4.**
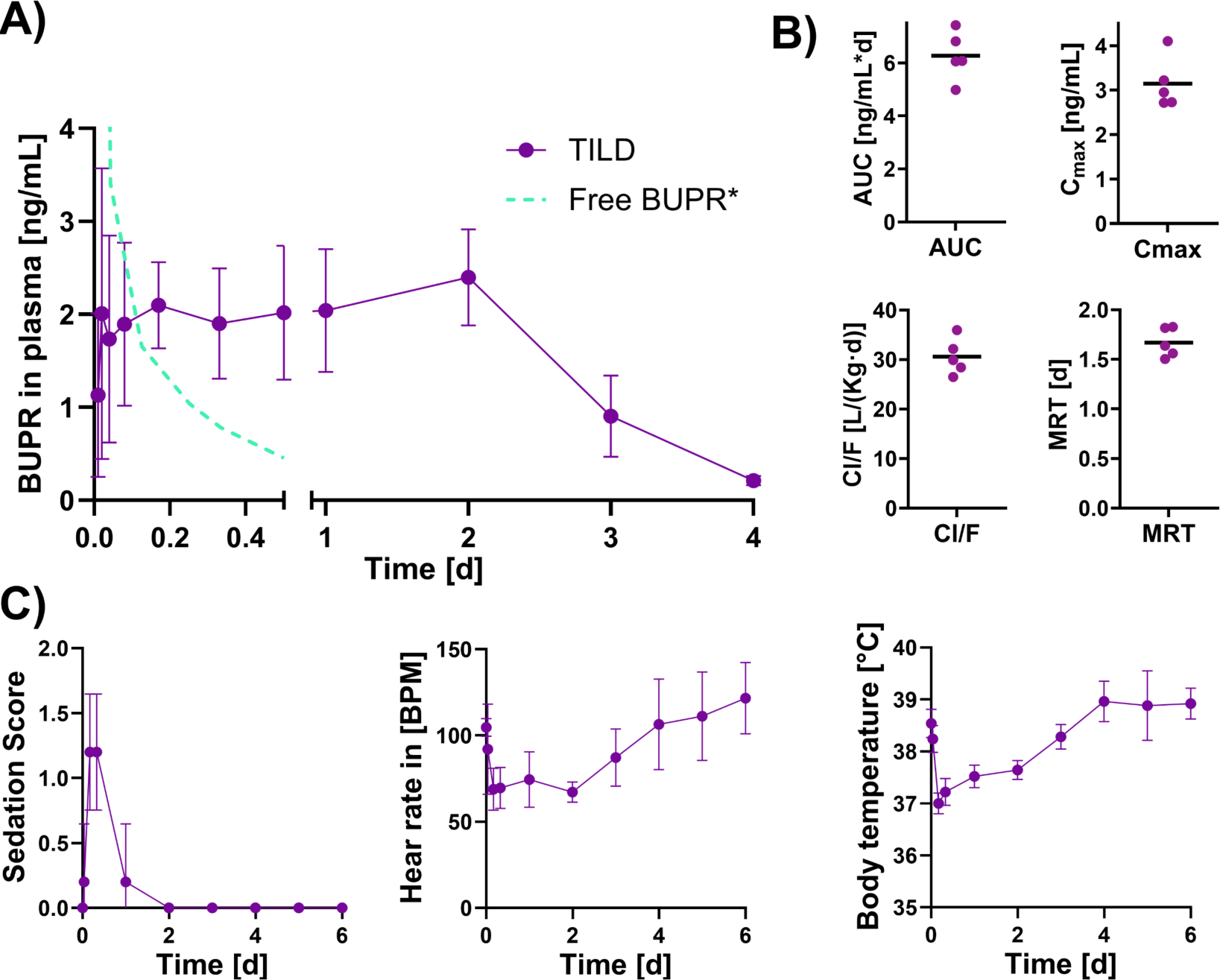
Pharmacokinetic and tolerability evaluation of TILD in beagle dogs (A) Plasma concentration-time profile of buprenorphine following subcutaneous administration of TILD compared to fast-acting buprenorphine (n=5; mean ±SD) (*data extrapolated from Nunamaker et al.[51]). (B) Pharmacokinetic parameters of TILD including AUC, C_max_, Clearance and MRT. (n=5; mean) (C) Evaluation of sedation scores and physiological parameters (vitality) following administration, indicating tolerability (See Supporting Information for details on scoring) (n=5; mean ±SD).

## Conclusion

In this study, we present TILD, a rationally designed liposomal drug delivery system that enables dual-mechanistic subcutaneous drug release via environmental responsiveness. Guided by computational simulations, we optimized the lipid composition to support both ion-triggered burst release and temperature-induced sustained diffusion, validated through SAXS, NMR, and DSC. TILD forms stable in situ depots driven by cation-mediated vesicle clustering, as confirmed by DLS and Cryo-TEM. These design features translated into prolonged pharmacokinetics, depot retention, and therapeutic efficacy in vivo. While pre-aggregated depot achieved comparable analgesic outcomes, TILD offers key advantages in terms of ease of formulation and injection, making it a more practical and scalable alternative to existing fast-acting or manually aggregated systems. Validation in rats confirmed pharmacodynamic efficacy of the formulation. A pharmacokinetic study in Beagle dogs further provided critical insights into translational readiness for veterinary and potentially human medicine. By integrating computational insights into formulation design, this study enabled the development of TILD - a scalable and translatable system for sustained analgesia with the potential to serve as a platform for broader application across injectable therapies.

## Materials and methods

### Materials

Phosphatidylcholine, hydrogenated (HSPC), 1,2-dioleoyl-*sn*-glycero-3-phospho-(1’-rac-glycerol) (sodium salt) (DOPG) 1,2-dioleoyl-*sn*-glycero-3-phosphocholine (DOPC); 1,2- distearoyl-*sn*-glycero-3-phosphocholine (DSPC) (Lipoid, Ludwigshafen, Germany); cholesterol (Chol), bromphenolblue and sodium chloride (NaCl) (Sigma Aldrich, Schnelldorf, Germany); Buprenorphine (BUP) was obtained from LGC Standards (Teddington, UK); Bupivacaine-d9 was obtained from Cayman Chemical Company, USA. LC/MS grade acetonitrile, water, and formic acid 98% − 100% and Nalaxone were purchased from Merck (Darmstadt, Germany); 1,1′-dioctadecyl-3,3,3′,3′-tetramethylindodicarbocyanine, 4-chlorobenzenesulfonate salt (DiD) (Invitrogen, Carlsbad, USA); 4-(2-hydroxyethyl)-1- piperazineethanesulfonic acid (HEPES), calcium chloride hexahydrate (CaCl_2_), magnesium chloride (MgCl_2_) hexahydrate (Carl Roth, Karlsruhe, Germany); isoflurane (Attane™, Piramal Pharma, India); pentobarbital (Esconarkon^®^; Streuli Tiergesundheit SA, Switzerland). Gibco™ water for injection (WFI) Fischer scientific (Switzerland). Acetonitrile (ACN), methanol (MeOH), chloroform, tetrahydrofuran (THF) and trifluoroacetic acid (TFA) were obtained from Carl Roth (Karlsruhe, Germany); Purified and deionized water (H_2_O) was prepared with Smart2Pure3 (Thermo Scientific, Niederelbert, Germany). Deuterated solvents such as D2O (99.9%) were obtained from Deutero GmbH (Kastelaun, Germany) and MeOD (99.80%) from Eurisotop (St.-Aubin Cedex, France).

### Methods

#### Molecular Dynamics Simulations

Atomistic molecular dynamics (MD) simulations were performed using GROMACS 2023.3 with lipid force fields generated through CHARMM-GUI v1.7. A bilayer membrane comprising HSPC and DOPG (75:25 mol%; Note: HSPC was composed as a mixture of DSPC at 67.5 mol% and DPPC at 7.5 mol%) was constructed in a simulation box of 12 × 12 × 10 nm. Buprenorphine molecules were incorporated at a 1:20 drug-to-lipid molar ratio. The system was solvated with water and physiological NaCl, and divalent cations were introduced to approximate and simulate subcutaneous (s.c.) conditions. Energy minimization was conducted using a steepest descent integrator for 1,000 steps with a 2 fs timestep (2ps). Equilibration followed by 2ns, and production simulations ran for 50ns to 100ns. All atomistic simulations were performed using the leap-frog integrator with a 2 fs time step over a total simulation time of 100 ns. Periodic boundary conditions (PBC) were applied in all directions (xyz). Electrostatic interactions were treated using the Particle Mesh Ewald (PME) method with a real-space cutoff (rcoulomb) of 1.2 nm, a PME order of 4, and a Fourier grid spacing of 0.16 nm. van der Waals interactions were modelled using a force-switch function (vdwtype = cutoff with vdw-modifier = force-switch), with a cutoff radius (rvdw) of 1.2 nm and switching beginning at 1.0 nm. Neighbour searching followed the Verlet cutoff scheme with a list radius of 1.2 nm. Temperature coupling was applied separately to solute and solvent groups using the V-rescale thermostat (tcoupl = V-rescale) with a time constant (tau_t) of 1.0 ps and a reference temperature of 311.15 K or temperature ramps from 338.15 K and 277.15 K. Semiisotropic pressure coupling was used via the C-rescale algorithm with tau_p = 5.0 ps, reference pressure set to 1.0 bar, and compressibility of 4.5×10⁻⁵ bar⁻¹ in the xy and z directions.[52] All bonds involving hydrogen atoms were constrained using the LINCS algorithm (lincs_order = 4).

Coarse-grained simulations were carried out using the Martini 3 force field.[56] The INSANE (insane.py; Version 1.2.0) script was used to generate the lipid bilayer starting configuration. The lipid bilayer made of 3360 lipids was inserted into a larger box with water and subsequently equilibrated. After approximately 80 ns of simulation, it spontaneously formed a liposome of approximately 22 nm diameter. Afterwards, two liposome replicas were placed into a simulation box of 35x35x55 nm with NaCl at physiological concentration of 150 mM. Two systems were simulated, one in which Ca^2+^ cations were added to the previously described system and another one without Ca^2+^. Each simulation was carried out for at least 3 µs. under isotropic pressure coupling. Simulations were conducted using the leap-frog integrator with a time step of 25 fs for a total of 3 μs. Periodic boundary conditions were applied in all spatial dimensions (xyz). Nonbonded interactions were computed using the Verlet cutoff scheme, with a cutoff distance of 1.1 nm for both van der Waals and Coulombic interactions. Electrostatics were treated using the reaction-field method (coulombtype = reaction-field) with a dielectric constant (epsilon_r) of 15 and reaction field dielectric (epsilon_rf) set to 0. van der Waals interactions were truncated at 1.1 nm using a simple cutoff scheme. Temperature coupling was applied separately to solvent and non-solvent groups using the V-rescale thermostat with a time constant (tau_t) of 4.0 ps and a reference temperature of 311 K. Isotropic pressure coupling was applied using the C-rescale barostat with a time constant (tau_p) of 4.0 ps, compressibility of 3×10⁻⁴ bar⁻¹, and a reference pressure of 1.0 bar. Neighbour lists were updated every 20 steps using a grid-based approach. All simulations were initialized with Maxwell-distributed velocities at 311 .15 K. Bond constraints were disabled (constraints = none), and LINCS was used for enforcing bond constraints during the simulation with lincs_order = 12 and lincs_iter = 2 for numerical stability. Simulation and trajectories analysis were performed using GROMACS 2023.3.

#### Liposome Preparation and Encapsulation Efficiency

Lipid films were prepared by evaporating chloroform solutions of lipids under nitrogen, followed by vacuum drying overnight. The dried film was hydrated with 25 mM HEPES, 140 mM NaCl buffer (pH 7.4) and subjected to six freeze–thaw cycles. Liposomes were then extruded ten times through stacked 200 nm polycarbonate membranes. Buprenorphine was loaded by passive encapsulation. Encapsulation efficiency (EE%) was determined using HPLC following separation by size exclusion chromatography (SEC) on Sephadex G50 columns. In brief, the HPLC mobile phase consisted of acetonitrile and PBS (83:17 v/v) with PBS prepared at 0.1 mM and pH adjusted to 6.0 using 1 M ortho-phosphoric acid, as adapted from literature.[57] For sample preparation, liposome samples were dissolved in a solvent mixture of mobile phase, and methanol (75:25 v/v), containing naloxone as an internal standard at a concentration of 0.1 mg/mL.

#### Small-angle X-ray Scattering (SAXS)

SAXS measurements were carried out to determine the atomic/molecular arrangement of lipids into liposomes. Measurements were performed on a Bruker AXS Micro, with a micro focused X-ray source, operating at voltage and filament current of 50 kV and 1000 μA, respectively. The Cu Kα radiation (λCu Kα = 1.5418 Å) was collimated by a 2D Kratky collimator, and the data were collected by a 2D Pilatus 100 K detector. The scattering vector Q = (4π/λ) sin θ, with 2θ being the scattering angle, was calibrated using silver behenate. Data were collected and azimuthally averaged using the Saxsgui software to yield 1D intensity vs. scattering vector Q, with a Q range from 0.001 to 0.5 Å–1. For all measurements, the samples were placed inside a quartz capillary cell with a sample volume of 100 μL and a thickness of ∼1,5 mm. Samples were equilibrated at 25 or 37 °C for 10 min prior to measurement, and the scattering intensity was collected for 2 h at 25 or 37 °C. Liposomes dispersion is irradiated with a narrow and collimated beam of quasi-monochromatic soft X-rays which interacts with the liposomal dispersion. The X-rays scattered by the vesicles are detected with a position-sensitive area detector. From a mathematical perspective, the recorded scattering pattern (Supporting Fig S7) is proportional to the square of the three-dimensional Fourier transform of the electron density distribution ρ(r) in the material, and thus, by analysing the SAXS profile, information on the nanoscale structure and morphology of the sample studied are quantified as already described in literature.[36]

#### Nuclear Magnetic Resonance (NMR)

For NMR analysis, liposomes were hydrated in D₂O-based HEPES buffer; buprenorphine was dissolved in CD₃OD as a reference. Liquid-state NMR of buprenorphine/CD_3_OD was performed on a Bruker Avance NEO spectrometer (500.13 MHz, 1.7 mm TXI triple-resonance, z-gradient) at 297 K. The ^1^H NMR spectrum was recorded using the *zg30* pulse program (Bruker pulse program library) applying a 30 degree flip angle, 64 scans, 1 s relaxation delay, 20 ppm spectral width, and 64 k data points.

High Resolution Magic Angle Spinning (HR-MAS) NMR of liposomal dispersions in 4 mm zirconium oxide MAS rotors with 50 µL Teflon inserts was carried out on a Bruker Avance II spectrometer (500.13 MHz, 4 mm HR-MAS dual ¹H/¹³C probe, magic angle gradient) at 293 K and–310 K and 8 kHz MAS rate. 1D ¹H spectra were acquired with the *zg* pulse sequence (Bruker pulse program library) applying 16–256 scans, 4 s relaxation delay, 16 ppm spectral width, and 32k data points. All data were processed in Bruker Topspin (v3.5 and 4.0.9).

#### Differential Scanning Calorimetry (DSC)

Thermal behaviour was analysed using a DSC 250 instrument (TA Instruments, USA). Multilamellar liposomes (50 mM lipid) were sealed in T_zero_^TM^ aluminium pans and scanned from −40 °C to 75 °C at a rate of 2 °C/min. Data from the second cycles were used for analysis of the thermal profile and hysteresis. Enthalpy and transition temperatures (Tm, onset) were calculated from the second heating cycle and normalized to lipid content using the TRIOS Software version 5.1.

#### Dynamic Light Scattering (DLS) and Zeta Potential

The hydrodynamic diameter, polydispersity index (PDI), and zeta potential were determined using a Litesizer 500 instrument (Anton Paar, Graz, Austria) operating with a 633 nm helium-neon laser at a 175° backscatter angle at 25 °C.[58] Samples were diluted in HEPES buffer (refractive index of 1.33) and measured with 30 runs of 10 s (size) and 100 runs (zeta potential) using the Smoluchowski approximation with a Debye factor of 1.5. The intensity-based size distribution of the liposomes was unimodal, allowing the autocorrelation function to be analysed using the cumulant method, as implemented in the Kalliope™ software.

#### Turbidity Measurements

Aggregation behaviour was assessed by measuring optical density at 400 nm (OD₄₀₀) in a quartz 96-well microtiter plate with a clear and flat bottom (Hellma GmbH & Co. KG, Germany). Liposomes were mixed 1:3 with divalent cation solutions of varying concentrations and incubated for 2 min before OD₄₀₀ was recorded using an Infinite plate reader (Tecan, Switzerland).

#### Cryogenic Transmission Electron Microscopy (Cryo-TEM)

Samples (4 µL of 1 mg/mL lipid) were applied to plasma-cleaned lacey carbon grids, blotted, and plunge-frozen in liquid ethane using a Vitrobot system under 100% humidity. Imaging was performed on a FEI Tecnai Spirit F20 TEM at 80 kV under low-dose conditions with an Eagle CCD camera.

#### In Vitro Release Studies

Drug release was evaluated using Franz (Side-by-Side) diffusion cells (2 mL volume per chamber, 9mm orifice diameter) with a 100 nm polycarbonate membrane separating the donor and acceptor chambers. TILD or control formulations were loaded into the donor chamber. The acceptor chamber was filled with HEPES buffer with or without added divalent cations and maintained at either 25 °C or 38 °C. At defined intervals, aliquots of 200 μL were withdrawn and analysed via HPLC. Drug release profiles were analysed by fitting both monophasic (first-order exponential) and biphasic (double exponential) models to the averaged experimental data. Model selection was performed by comparing the coefficient of determination (R2), values to further assess the goodness-of-fit and Akaike Information Criterion (AIC), and Bayesian Information Criterion (BIC) comparison to distinguish between the two kinetic models. Analysis were performed using Python (version 3.13. 3).

#### In vivo experiments

All animal experiments were performed on 12-week-old Lewis rats (BW: F185g / M330g) (LEW/OrlRj, Janvier Laboratories, France) in accordance with institutional and federal regulations governing animal care and use, according to the ARRIVE guidelines, and were approved by the Cantonal Veterinary Office of Bern (Switzerland) (BE47/2021). Rats were housed in specific pathogen-free (SPF) conditions in compliance with the and Federation of European Laboratory Animal Science Associations (FELASA) guidelines. Up to 3 rats were be accommodated in autoclaved IVCs cages (Blue line, Tecniplast, Italy) with aspen wood bedding (J. Rettenmeier & Söhne GmbH, Germany) and enrichments. The housing conditions were controlled with a 12:12 h light: dark cycle, room temperature in the range of 22+/-2 °C and relative humidity in the range of 45%-65%. Autoclaved tap water and irradiated rodent chow (Mouse and Rat Maintenance 3432, Granovit, Switzerland) were provided ad libitumin. The rats underwent a one-week acclimatization period and were regularly handled by personnel for gentling and habituation to the procedures. On the day of experiment, animals were randomly allocated to different treatment groups, and treatments were blinded for the researchers. Each group, consisting of equivalent males and females, received a different treatment and the sex was then imposed as blocking factor for the statistical analysis. The results are presented as mean ± σ or mean ± SEM .

#### Biodistribution

Twelve-week-old Lewis rats (n = 4/group, sex balanced) were anesthetized with 5% isoflurane in 100% O₂ at 1 L/min for induction. Anaesthesia was maintained at 2% minimum alveolar concentration via a face mask. Each animal received a subcutaneous injection of 150 μL of DiD-labeled liposome formulation (100 mM lipid concentration, 0.01 mol% DiD) into the shaved back of the neck. Fluorescence imaging was performed using the IVIS Spectrum CT scanner (PerkinElmer). After the final time point, animals were euthanized with intraperitoneal injection of pentobarbital (150 mg/kg), and major organs were harvested to assess systemic biodistribution. The IVIS scanner was operated in fluorescence imaging mode with an emission filter set at 650 nm. Imaging conditions were standardized: a subject height of 1.5 cm, chamber temperature of 37 °C, and exposure time of 0.2 s. Image analysis was conducted using a fixed fluorescence scale (1’000–40’000 counts). Regions of interest (ROIs) were manually defined for each injection site and kept consistent across all images. Total Fluorescence Counts (TFC, in photons/second) were calculated by summing all pixel values within each ROI to assess local retention and clearance dynamics.

#### In Vivo Pharmacokinetics

Twelve-week-old Lewis rats (n = 6/group, sex balanced) received 0.1 mg/kg free drug (Temgesic injection) or 1.2 mg/kg of buprenorphine encapsulated into the TILD system via subcutaneous injection (n=6/group, sex-balanced) at the base of the neck. Blood samples (100 μL) were collected from the lateral tail vein at the following time points: −2 days (baseline), 0.25, 2, and 8 h, and 1, 2, 3, 4, 7, 9, 11, and 14 days post-injection. Sampling was performed with conscious animals placed in a restrainer. Animals were euthanized by CO₂ inhalation after the final sampling point. Collected blood was transferred into K₂-EDTA BD Microtainer™ tubes (Fisher Scientific AG, Switzerland), centrifuged at 4 °C for 5 min at 3’000 g, and the resulting plasma (25 μL) was stored at −20 °C until analysis. For drug quantification, buprenorphine (BUP) was extracted from plasma by adding 125 μL of methanol containing deuterated buprenorphine as an internal standard and 100 μL of acetonitrile. Samples were centrifuged at 20’000 g for 1 hour at 4 °C, and the drug concentration in the resulting supernatant was measured using LC-MS/MS on a Sciex QTrap 5500 instrument (AB Sciex Switzerland GmbH) equipped with an UltiMate 3000 HPLC system (Thermo Fisher Scientific, Switzerland). For quantitation of BUP by multiple reaction monitoring the precursor/product ion transition m/z 468.3/396.3 was used as the quantifier and the transition m/z 468.3/413.9 served as the qualifier. Buprenorphine-d4 (Merck, Switzerland) was used as the internal standard (m/z 472.3 /400.3).

#### In Vivo Pharmacodynamics

Postoperative analgesic efficacy was evaluated in a rat plantar incision model using twelve-week-old Lewis rats (n = 12/group, sex balanced).[59] In brief, the plantar aspect of one hind paw was prepared using a 10% povidone-iodine solution in a sterile setting. The foot was placed through an opening in a sterile drape. Formulations were administered subcutaneously at the base of the neck at a dose of 1.2 mg/kg prior to surgery. A 1-cm longitudinal incision was made on the plantar surface of the paw using a No. 11 scalpel blade, beginning approximately 0.5 cm from the proximal edge of the heel and extending toward the toes. The underlying plantaris muscle was elevated and incised longitudinally, while maintaining the muscle’s origin and insertion. Hemostasis was achieved using gentle pressure. The skin was closed with two mattress sutures (5-0 nylon, FS-2 needle). Postoperatively, incisions were monitored daily for signs of infection, dehiscence, or hematoma. Animals presenting any such conditions were excluded from further analysis. After recovery from anesthesia, unrestrained rats were placed in clear plastic chambers with elevated mesh floors and allowed to acclimate until they were resting undisturbed. Mechanical withdrawal thresholds were assessed using electronic von Frey filaments (Bioseb, France), applied from below through the mesh to the plantar surface adjacent to the incision (typically medial to the incision near the heel). The contralateral, unoperated paw served as an internal control. Three measurements per paw were taken at each time point for each animal. The average value was normalized to the individual pre-surgical baseline measurement for comparison of treatment effects. To determine if there were statistically significant differences between the TILD and Pre-formed Depot groups across time, a linear mixed-effects model (LME) was used. The LME was selected to account for the repeated measurements taken from individual animals over time (random effect) and to handle the unbalanced nature of the dataset. Fixed effects included Group, Time, and their interaction.

#### Pharmacokinetic and tolerability evaluation

To assess the pharmacokinetic profile and tolerability of TILD in a large-animal model, a study was conducted in five non-naïve male Beagle dogs (aged 8–48 months). Animals were under controlled conditions, including a two-week washout period prior to dosing. The animals received a single subcutaneous administration of TILD at a buprenorphine dose of 0.2 mg/kg. Blood samples for pharmacokinetic analysis were collected in EDTA K2 or K3 tubes via catheter at the following timepoints: pre-dose, 0.25, 0.5, 1, 2, 4, 8, and 12 h, and daily up to 6 days post-administration. Plasma was separated and stored at –80 °C in 500 µL aliquots for later analysis by LC/MS. Further, animals were closely monitored for local and systemic safety outcomes throughout the study. Vital parameters - including heart rate, respiratory rate, oxygen saturation, and body temperature - were assessed at baseline, once between 4–6 h post-dose, and once daily for the following six days. Signs of gastrointestinal distress, such as vomiting, obstipation or diarrhea, were recorded. Local dermal reactions at the injection site were scored and photo-documented at the same time intervals. Behavioral and general health observations, including videos (24 h post-injection), were made during dosing and blood collection periods for supplementary behavioral assessment. Safety experiments in Beagle dogs were conducted at the Institut National de la Recherche Scientifique (INRS), Canada, under AAALAC and CCAC certification.

### Statistics and reproducibility

All statistical analyses performed are described in the corresponding figure legends and within the Results and Discussion sections of the manuscript. GraphPad Prism 10.2.3 (GraphPad Software, USA), R version 4.4.0 (The R Foundation, Germany) and Python version 3.13. 3 were used for statistical evaluations. Group comparisons were made using appropriate parametric or non-parametric tests as specified. All experiments were conducted with independent biological replicates, and no data were excluded from the analysis. Animal studies were performed in a randomized and blinded manner, with sex included as a blocking factor to control for biological variability. Sample sizes and replicates were chosen based on previous experience with similar systems.

## Data availability

All data generated or analyzed during this study are included in this published article and its supplementary information file. The files to run MD simulations are publicly available on GitHub (https://github.com/Luciani-Group/MD_Liposome_SC).

## Author contributions

RE: Conceptualization, Methodology, Software, Investigation, Validation, Data curation, Visualization, Formal analysis, Writing – original manuscript.

SA: Conceptualization, Methodology, Investigation, Supervision, Data curation, Writing – Review & editing

JB: Software, Investigation, Validation, Formal analysis, Visualization, Writing – Review & editing

DB: Methodology, Software, Supervision, Writing – Review & editing LB: Methodology, Investigation, Visualization, Formal analysis

BH: Methodology, Formal analysis, Writing – Review & editing SS: Methodology, Investigation

MV: Methodology, Investigation, Validation, Formal analysis PV: Methodology, Investigation, Validation, Formal analysis

GR: Conceptualization, Methodology, Software, Supervision, Funding acquisition, Resources, Writing – Review & editing

AB: Conceptualization, Methodology, Funding acquisition, Supervision, Resources, Writing – Review & editing

PL: Conceptualization, Supervision, Project administration, Funding acquisition, Resources, Writing – Review & editing.

## Acknowledgement

We thank Claudia Spadavecchia and Olivier Levionnois for constructive feedback on the veterinary aspect of the study, and Unitectra for financial support of the in vivo study in Beagle dogs. Further, we thank Kevin Weber-Wilk for support during PK studies, Maria Nanni and Carlotta Lambertini for guidance and surgical training. The authors thank Giorgio Buttitta for the support with formulation preparation and Elia Wasserfallen for evaluating production scalability. Also, we would like to thank Claudia Bühr for technical assistance with LC/MS analytics. The authors would further like to thank Raffaele Mezzenga for his generous support on the SAXS measurements.

Figures created with Biorender.

## References

[1] A.M. Vargason, A.C. Anselmo, S. Mitragotri, The evolution of commercial drug delivery technologies, Nat Biomed Eng 5 (2021) 951–967. 10.1038/s41551-021-00698-w.

[2] V.R. Feig, S. Park, P.G. Rivano, J. Kim, B. Muller, A. Patel, C. Dial, S. Gonzalez, H. Carlisle, F. Codreanu, A. Lopes, A.E. Erdogan, N. Fabian, A. Guevara, A. Pettinari, J. Li, J. Liang, G.W. Liu, M.W. Tibbitt, G. Traverso, Self-aggregating long-acting injectable microcrystals, Nat Chem Eng 2 (2025) 209–219. 10.1038/s44286-025-00194-x.

[3] Wei Li, Jie Tang, Dennis Lee, Thomas R. Tice, Steven P. Schwendeman, Mark R. Prausnitz, Clinical translation of long-acting drug delivery formulations, Nat Rev Mater 7 (2022) 406–420. 10.1038/s41578-021-00405-w.

[4] M.J. Kreek, B. Reed, E.R. Butelman, Current status of opioid addiction treatment and related preclinical research, Sci Adv 5 (2019) eaax9140. 10.1126/sciadv.aax9140.

[5] C. Wang, X. Liu, W. Lv, X. Kuang, F. Wu, X. Fan, Y. Pang, Long-lasting comfort ocular surface drug delivery by in situ formation of an adhesive lubricative Janus nanocoating, Sci Adv 11 (2025) eads0282. 10.1126/sciadv.ads0282.

[6] C.M. Manasa, U. Likhitha, U.Y. Nayak, Revolutionizing animal health: A comprehensive review of long-acting formulations, J Drug Deliv Sci Technol 101 (2024). 10.1016/j.jddst.2024.106226.

[7] J.M. da Costa, T.G. Barroso, J.C. Prata, Research priorities in veterinary palliative care, Vet J 306 (2024) 106184. 10.1016/j.tvjl.2024.106184.

[8] D.S. Goldberg, S.J. McGee, Pain as a global public health priority, BMC Public Health 11 (2011). 10.1186/1471-2458-11-770.

[9] N. Lintzeris, A.J. Dunlop, P.S. Haber, D.I. Lubman, R. Graham, S. Hutchinson, S. Arunogiri, V. Hayes, P. Hjelmström, A. Svedberg, S. Peterson, F. Tiberg, Patient-Reported Outcomes of Treatment of Opioid Dependence With Weekly and Monthly Subcutaneous Depot vs Daily Sublingual Buprenorphine A Randomized Clinical Trial, JAMA Netw Open 4 (2021) E219041. 10.1001/jamanetworkopen.2021.9041.

[10] Tomasini Lorenzo, Ferrere Marianne, Nicolas Julien, Subcutaneous drug delivery from nanoscale systems, Nat Rev Bioeng 2 (2024) 501–520. 10.1038/s44222-024-00161-w.

[11] R. Sharma, S. Yadav, V. Yadav, J. Akhtar, O. Katari, K. Kuche, S. Jain, Recent advances in lipid-based long-acting injectable depot formulations, Adv Drug Deliv Rev 199 (2023). 10.1016/j.addr.2023.114901.

[12] J. Li, Z. Xia, M. Yu, A. Schwendeman, Challenges in the development of long acting injectable multivesicular liposomes (DepoFoam® technology), EJPB 205 (2024). 10.1016/j.ejpb.2024.114577.

[13] R. Eugster, P. Luciani, Liposomes: Bridging the gap from lab to pharmaceuticals, Curr Opin Colloid Interface Sci 75 (2025) 101875. 10.1016/j.cocis.2024.101875.

[14] M.N. O’Brien, W. Jiang, Y. Wang, D.M. Loffredo, Challenges and opportunities in the development of complex generic long-acting injectable drug products, J Con Rel 336 (2021) 144–158. 10.1016/j.jconrel.2021.06.017.

[15] S. Alidori, R. Subramanian, R. Holm, Patient-Centric Long-Acting Injectable and Implantable Platforms─An Industrial Perspective, Mol Pharm 21 (2024) 4238–4258. 10.1021/acs.molpharmaceut.4c00665.

[16] H. Koppisetti, S. Abdella, D.D. Nakmode, F. Abid, F. Afinjuomo, S. Kim, Y. Song, S. Garg, Unveiling the Future: Opportunities in Long-Acting Injectable Drug Development for Veterinary Care, Pharmaceutics 17 (2025) 626. 10.3390/pharmaceutics17050626.

[17] L. Rahnfeld, J. Thamm, F. Steiniger, P. van Hoogevest, P. Luciani, Study on the in situ aggregation of liposomes with negatively charged phospholipids for use as injectable depot formulation, Colloids Surf B Biointerfaces 168 (2018) 10–17. 10.1016/j.colsurfb.2018.02.023.

[18] S. Aleandri, L. Rahnfeld, D. Chatzikleanthous, A. Bergadano, C. Bühr, C. Detotto, S. Fuochi, K. Weber-Wilk, S. Schürch, P. van Hoogevest, P. Luciani, Development and in vivo validation of phospholipid-based depots for the sustained release of bupivacaine, Eur J Pharm Biopharm 181 (2022) 300–309. 10.1016/j.ejpb.2022.11.019.

[19] C. Wang, X. Lan, L. Zhu, Y. Wang, X. Gao, J. Li, H. Tian, Z. Liang, W. Xu, Construction Strategy of Functionalized Liposomes and Multidimensional Application, Small 20 (2024). 10.1002/smll.202309031.

[20] T. Duran, A. P. Costa, J. Kneski, X. Xu, D. J. Burgess, H. Mohammadiarani, B. Chaudhuri, Manufacturing process of liposomal Formation: A coarse-grained molecular dynamics simulation, Int J Pharm 659 (2024) 124288. 10.1016/j.ijpharm.2024.124288.

[21] S. Davoudi, A. Ghysels, Defining permeability of curved membranes in molecular dynamics simulations, Biophys J 122 (2023) 2082–2091. 10.1016/j.bpj.2022.11.028.

[22] R. Eugster, M. Orsi, G. Buttitta, N. Serafini, M. Tiboni, L. Casettari, J.-L. Reymond, S. Aleandri, P. Luciani, Leveraging machine learning to streamline the development of liposomal drug delivery systems, J Con Rel 376 (2024) 1025–1038. 10.1016/j.jconrel.2024.10.065.

[23] L.H. Chen, J.N. Hu, Development of nano-delivery systems for loaded bioactive compounds: using molecular dynamics simulations, Crit Rev Food Sci Nutr (2024) 1811–1832. 10.1080/10408398.2023.2301427.

[24] M. Hale, M. Garofoli, R.B. Raffa, Benefit-risk analysis of buprenorphine for pain management, J Pain Res 14 (2021) 1359–1369. 10.2147/JPR.S305146.

[25] T.P. Clark, The history and pharmacology of buprenorphine: New advances in cats, J Vet Pharmacol Ther 45 (2022) S1–S30. 10.1111/jvp.13073.

[26] S. Spinella, R. McCarthy, Buprenorphine for Pain: A Narrative Review and Practical Applications, Am J Med 137 (2024) 406–413. 10.1016/j.amjmed.2024.01.022.

[27] S.S.C. Wong, T.H. Chan, F. Wang, T.C.W. Chan, H.C. Ho, C.W. Cheung, Analgesic Effect of Buprenorphine for Chronic Noncancer Pain: A Systematic Review and Meta-analysis of Randomized Controlled Trials, Anesth Analg 137 (2023) 59–71. 10.1213/ANE.0000000000006467.

[28] C. Timko, M.C. Lor, S. Kertesz, K. Kroenke, K. Macia, A. Nevedal, K.J. Hoggatt, Management of patients at risk of harms from both continuing and discontinuing their long-term opioid therapy: A qualitative study to inform the gap in clinical practice guidelines, Pain Practice (2024). 10.1111/papr.13440.

[29] R. Downing, G. Della Rocca, Pain in Pets: Beyond Physiology, Animals 13 (2023). 10.3390/ani13030355.

[30] J. Parchekani, A. Allahverdi, M. Taghdir, H. Naderi-Manesh, Design and simulation of the liposomal model by using a coarse-grained molecular dynamics approach towards drug delivery goals, Sci Rep 12 (2022) 2371. 10.1038/s41598-022-06380-8.

[31] H.M. Kinnunen, R.J. Mrsny, Improving the outcomes of biopharmaceutical delivery via the subcutaneous route by understanding the chemical, physical and physiological properties of the subcutaneous injection site, J Con Rel 182 (2014) 22–32. 10.1016/j.jconrel.2014.03.011.

[32] I. Torres-Terán, M. Venczel, S. Klein, Prediction of subcutaneous drug absorption − Development of novel simulated interstitial fluid media for predictive subcutaneous in vitro assays, Int J Pharm 658 (2024). 10.1016/j.ijpharm.2024.124227.

[33] H.M. Kinnunen, R.J. Mrsny, Improving the outcomes of biopharmaceutical delivery via the subcutaneous route by understanding the chemical, physical and physiological properties of the subcutaneous injection site, J Con Rel 182 (2014) 22–32. 10.1016/j.jconrel.2014.03.011.

[34] P. Hamal, V. Subasinghege Don, H. Nguyenhuu, J.C. Ranasinghe, J.A. Nauman, R.L. McCarley, R. Kumar, L.H. Haber, Influence of Temperature on Molecular Adsorption and Transport at Liposome Surfaces Studied by Molecular Dynamics Simulations and Second Harmonic Generation Spectroscopy, J Phys Chem B 125 (2021) 10506–10513. 10.1021/acs.jpcb.1c04263.

[35] C. Doyen, E. Larquet, P.D. Coureux, O. Frances, F. Herman, S. Sablé, J.P. Burnouf, C. Sizun, E. Lescop, Nuclear Magnetic Resonance Spectroscopy: A Multifaceted Toolbox to Probe Structure, Dynamics, Interactions, and Real-TimeIn SituRelease Kinetics in Peptide-Liposome Formulations, Mol Pharm 18 (2021) 2521–2539. 10.1021/acs.molpharmaceut.1c00037.

[36] L. Baraldi, S.R. Alfarano, F. Sousa, R. Mezzenga, B. Rothen-Rutishaser, A. Petri-Fink, S. Balog, Describing the SAXS Profile of Unilamellar Liposomes via Beating Waves, ChemRxiv (2023). 10.26434/chemrxiv-2023-nsbtp.

[37] C. Doyen, E. Larquet, P.D. Coureux, O. Frances, F. Herman, S. Sablé, J.P. Burnouf, C. Sizun, E. Lescop, Nuclear Magnetic Resonance Spectroscopy: A Multifaceted Toolbox to Probe Structure, Dynamics, Interactions, and Real-TimeIn SituRelease Kinetics in Peptide-Liposome Formulations, Mol Pharm 18 (2021) 2521–2539. 10.1021/acs.molpharmaceut.1c00037.

[38] M. Regenold, K. Kaneko, X. Wang, H.B. Peng, J.C. Evans, P. Bannigan, C. Allen, Triggered release from thermosensitive liposomes improves tumor targeting of vinorelbine, J Con Rel 354 (2023) 19–33. 10.1016/j.jconrel.2022.12.010.

[39] G. Buttitta, S. Bonacorsi, C. Barbarito, M. Moliterno, S. Pompei, G. Saito, I. Oddone, G. Verdone, D. Secci, S. Raimondi, Scalable microfluidic method for tunable liposomal production by a design of experiment approach, Int J Pharm 662 (2024). 10.1016/j.ijpharm.2024.124460.

[40] R. Sarabia-Estrada, A. Cowan, B.M. Tyler, M. Guarnieri, Association of nausea with buprenorphine analgesia for rats, Lab Anim (NY) 46 (2017) 242–244. 10.1038/laban.1277.

[41] A. Wolter, C.H. Bucher, S. Kurmies, V. Schreiner, F. Konietschke, K. Hohlbaum, R. Klopfleisch, M. Löhning, C. Thöne-Reineke, F. Buttgereit, J. Huwyler, P. Jirkof, A.E. Rapp, A. Lang, A buprenorphine depot formulation provides effective sustained post-surgical analgesia for 72 h in mouse femoral fracture models, Sci Rep 13 (2023). 10.1038/s41598-023-30641-9.

[42] A. Burton, D.J. DeBona, M. Handzel, S. Kelly-Pisciotti, M. Qiao, D. Rojek, N.M. Acquisto, Surgical removal of extended-release buprenorphine depot due to adverse reactions, Am J Emerg Med 81 (2024) 127–128. 10.1016/j.ajem.2024.04.047.

[43] T.J. Brennan, Postoperative Models of Nociception, ILAR J 40 (1999) 129–136. https://academic.oup.com/ilarjournal/article/40/3/129/959762.

[44] L. Carbone, Ethical and IACUC Considerations Regarding Analgesia and Pain Management in Laboratory Rodents, Comp Med 69 (2019) 443–450. 10.30802/AALAS-CM-18-000149.

[45] E. Laboureyras, J. Chateauraynaud, P. Richebé, G. Simonnet, Long-term pain vulnerability after surgery in rats: Prevention by nefopam, an analgesic with antihyperalgesic properties, Anesth Analg 109 (2009) 623–631. 10.1213/ane.0b013e3181aa956b.

[46] Leslie I Curtin, Julie A Grakowsky, Mauricio Suarez, Alexis C Thompson, Jean M DiPirro, Lisa BE Martin, Mark B Kristal, Evaluation of Buprenorphine in a Postoperative Pain Model in Rats, Comp Med 59 (2009) 60–71.

[47] M., Brennan, A., Sinusas, T. Horvath, Correlation between body weight changes and postoperative pain in rats treated with meloxicam or buprenorphine, Lab Anim (NY) 38 (2009) 87–93. 10.1038/laban0309-87.

[48] C.J. Rini, B.C. Roberts, A. Vaidyanathan, A. Li, R. Klug, D.B. Sherman, R.J. Pettis, Enabling faster subcutaneous delivery of larger volume, high viscosity fluids, Expert Opin Drug Deliv 19 (2022) 1165–1176. 10.1080/17425247.2022.2116425.

[49] D.H. Dyson, Perioperative Pain Management in Veterinary Patients, Veterinary Clinics of North America - Small Animal Practice 38 (2008) 1309–1327. 10.1016/j.cvsm.2008.06.006.

[50] Harshitha Shravya, Trilokchandran, Vijaykumar, Beagles in Biomedical Research: Scientific Justification, Biosafety Protocols, Ethical Debates, and Emerging Alternatives, IRE Journal 8 (2025) 1020–1026.

[51] Elizabeth A Nunamaker, DeAnne F Stolarik, Junli Ma, Amanda S Wilsey, Gary J Jenkins, Chris L Medina, Clinical Efficacy of Sustained-Release Buprenorphine with Meloxicam for Postoperative Analgesia in Beagle Dogs Undergoing Ovariohysterectomy, Journal of the American Association for Laboratory Animal Science 53 (2014) 494–501.

[52] G. Sinha, Companion therapeutics, Nat Biotechnol 32 (2014) 12–14. 10.1038/nbt.2793.

[53] C. Abboud, A. Duveau, R. Bouali-Benazzouz, K. Massé, J. Mattar, L. Brochoire, P. Fossat, E. Boué-Grabot, W. Hleihel, M. Landry, Animal models of pain: Diversity and benefits, J Neurosci Methods 348 (2021). 10.1016/j.jneumeth.2020.108997.

[54] K.E. Sadler, J.S. Mogil, C.L. Stucky, Innovations and advances in modelling and measuring pain in animals, Nat Rev Neurosci 23 (2022) 70–85. 10.1038/s41583-021-00536-7.

[55] Amir Kol, Boaz Arzi, Kyriacos A. Athanasiou, Diana L. Farmer, Jan A. Nolta, Robert B. Rebhun, Xinbin Chen, Leigh G. Griffiths, Frank J. M. Verstraete, Christopher J. Murphy, Dori L. Borjesson, Companion animals: Translational scientist’s new best friends, Sci Transl Med 7 (2015) 308ps21. 10.1126/scitranslmed.3003276.

[56] P.C.T. Souza, R. Alessandri, J. Barnoud, S. Thallmair, I. Faustino, F. Grünewald, I. Patmanidis, H. Abdizadeh, B.M.H. Bruininks, T.A. Wassenaar, P.C. Kroon, J. Melcr, V. Nieto, V. Corradi, H.M. Khan, J. Domański, M. Javanainen, H. Martinez-Seara, N. Reuter, R.B. Best, I. Vattulainen, L. Monticelli, X. Periole, D.P. Tieleman, A.H. de Vries, S.J. Marrink, Martini 3: a general purpose force field for coarse-grained molecular dynamics, Nat Methods 18 (2021) 382–388. 10.1038/s41592-021-01098-3.

[57] D.A. Mundhey, V. V. Rajkondawar, A.S. Daud, N.P. Sapkal, A study of method development, validation and forced degradation for quantification of buprenorphine hydrochloride in a microemulsion formulation, J Appl Pharm Sci 6 (2016) 159–169. 10.7324/JAPS.2016.601022.

[58] R.T. Coones, I. Nikolic, R. Eugster, D. Mehn, V. Tong, P. Luciani, C. Minelli, Best practice for the size analysis of nanomedicines – An iron sucrose case study, Int J Pharm 674 (2025). 10.1016/j.ijpharm.2025.125452.

[59] T.J. Brennan, E.P. Vandermeulen, G.F. Gebhart, Characterization of a rat model of incisional pain, Pain 64 (1996) 493–501. 10.1016/0304-3959(95)01441-1.

