## Supporting Information for "Hot cue: Physiologically controlled release from an in situ forming liposomal depot"

*Molecular Dynamics Simulations*

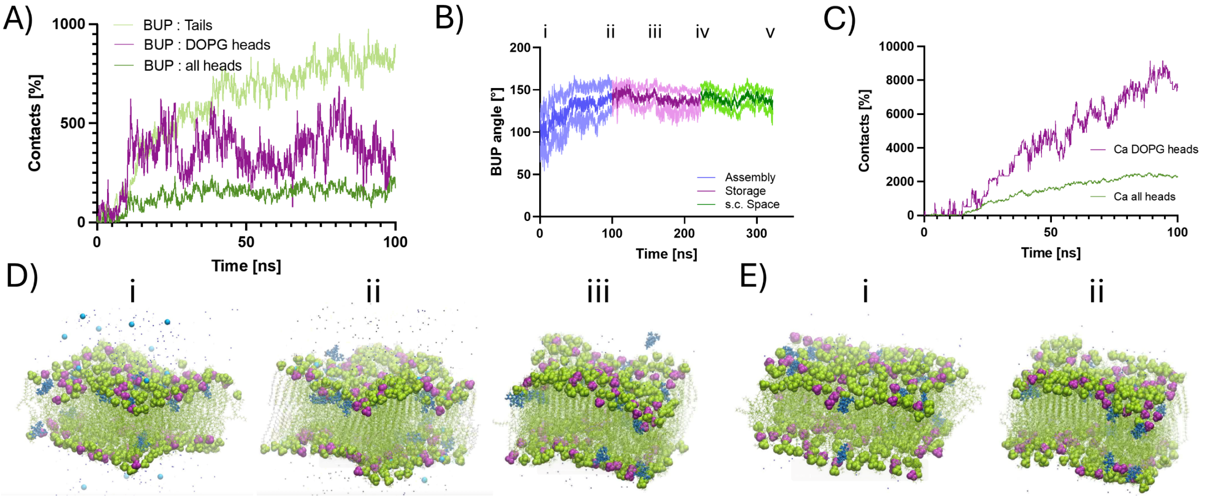

Fig S1 (A) Contacts between drug molecules and various lipid components, showing that drug molecules predominantly position themselves within the lipid bilayer and interact with the negatively charged headgroups of DOPG during drug loading and constructing the system. (B) Orientation of the drug molecule within the lipid bilayer over consecutive simulations i. start of drug loading (65 °C); ii. Drug incorporated into membrane iii. Cool down to storage temperature (4 °C); iv. Warm up to body temperature and exposure to simulated s.c. space; v. After exposition and surface associated drug release. (C) Number of contacts between divalent cations (Ca) and lipid headgroups, indicating a preferential interaction with DOPG headgroups, confirming competitive binding. (D) Snapshots of membrane simulation in approximated s.c. environment i) Simulation at 0ns with divalent cations dispersed in the simulation box and drug molecules associated with the membrane ii) 50ns showing divalent cations have moved towards the membrane and drug molecules start to decouple from the membrane surface iii) 100ns drug molecules fully dissociated from the membrane. E) Snapshots of membrane across temperature ramp i) Fluid membrane and ii) rigid gel membrane.

*Nuclear magnetic resonance measurements*

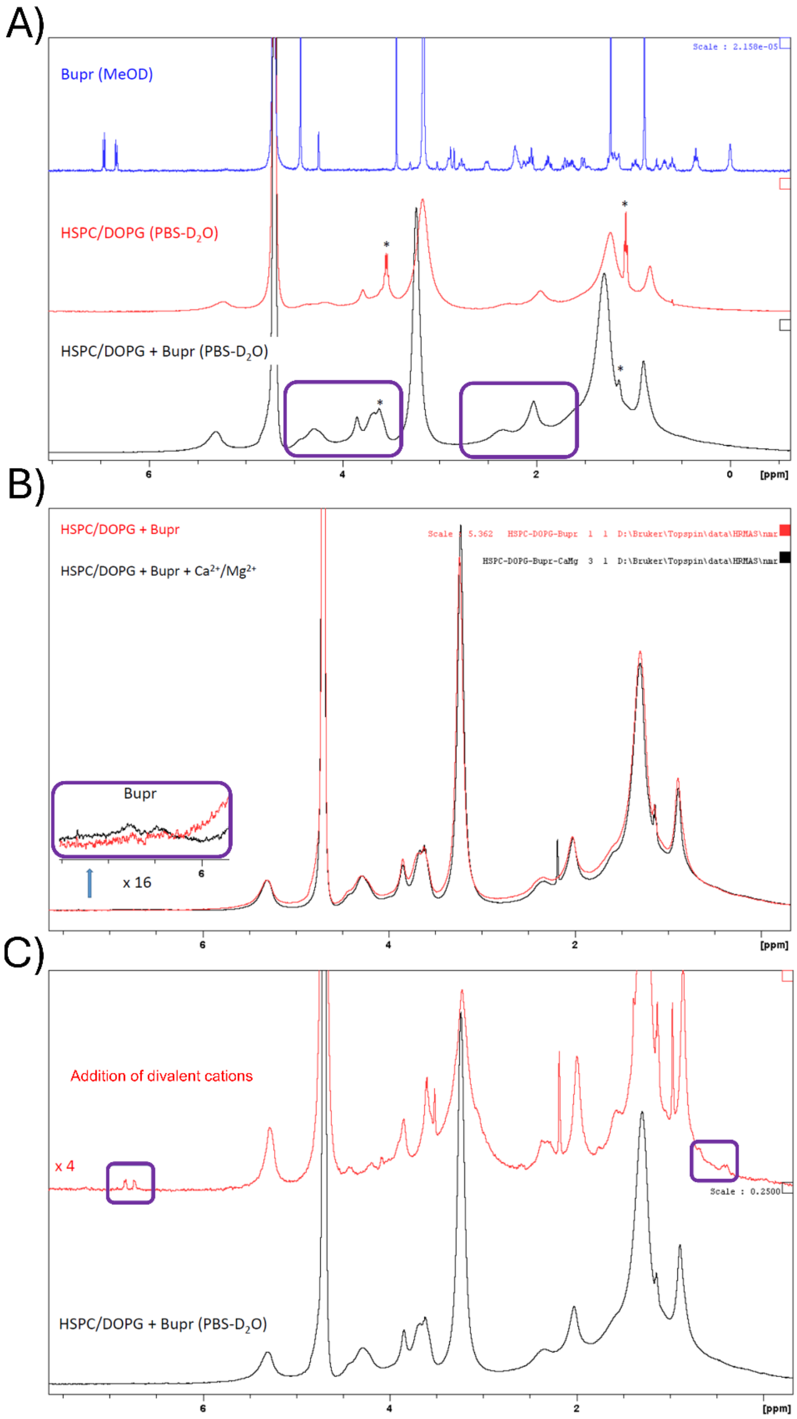

Fig S2 Liquid state $^{1}$H NMR (Bupr/MeOD) and HR-MAS $^{1}$H NMR spectra of liposomes under varying conditions (T = 293 K). A) Spectra of free buprenorphine (blue), empty liposomes composed of DOPG:HSPC (25:75 mol%) (red), and buprenorphine-loaded liposomes (black). B) Spectra of buprenorphine-loaded liposomes before and after exposure to divalent cations at physiological concentrations, representative of the subcutaneous (s.c.) environment. C) Spectra of drug-loaded vesicles following further addition of divalent cations, illustrating enhanced drug dissociation from the vesicle surface. Purple boxes highlight resonances from buprenorphine. Peaks marked with * derive from ethanol impurity.

*Differential Scanning Calorimetry*

Table S1 Phase transition temperature of lipid mixtures (50 mM) with drug molecule incorporated.

| Formulation | Enthalpy J/g | Onset °C |
|---|---|---|
| 1. HSPC:DOPG (75:25) | 38.45 | 34.38 |
| 2. HSPC:DOPG (80:20) | 37.83 | 39.98 |
| 3. HSPC:DOPG (70:30) | 27.02 | 32.14 |
| 4. HSPC:DOPG (75:25) + 100 mM Zn | 33.06 | 42.99 |
| 5. HSPC:DOPC (75:25) | 48.79 | 40.71 |

*Aggregation Studies*

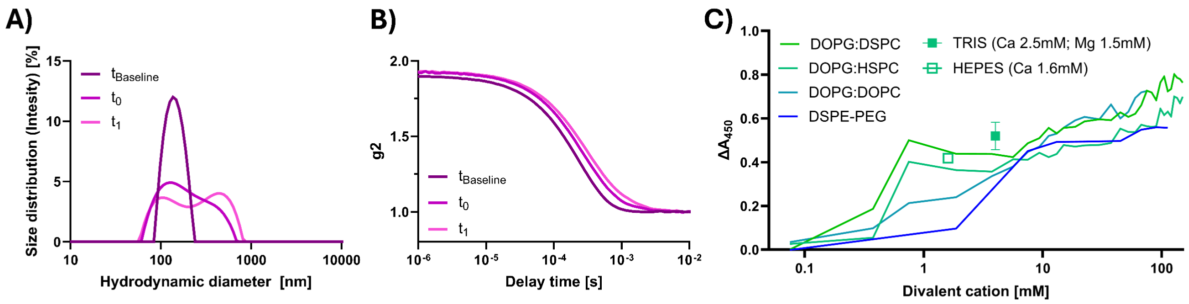

Fig S3 Aggregation studies of liposomes A) Size distribution and B) auto correlation function of liposomes exposed to physiological cation concentrations over time indicating aggregation behavior C) Aggregation profiles of different lipid systems along an increasing concentration gradient of divalent cations. TRIS & HEPES, represent two points simulating the physiological conditions in the s.c. space where the lead formulation (DOPG:HSPC 25:75 n%) was additionally screened. (Note: all lipid compositions were 25:75 n%; DSPE-PEG= DOPG:HSPC:DSPE-PEG (25:73.5:1.5 n%))

*Stability studies*

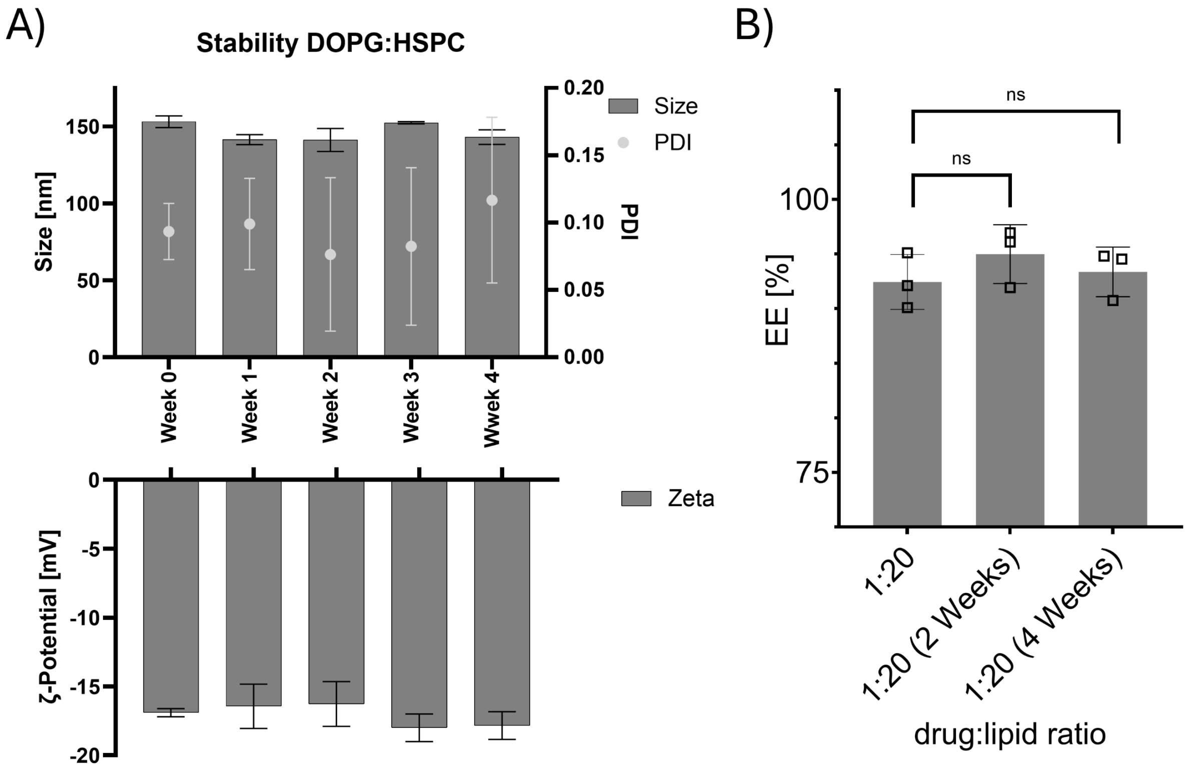

Fig S4 Drug delivery system (DOPG:HSPC 25:75 n%) stability over 4 weeks considering A) Size and zeta potential and B) drug molecule encapsulation (mean ± SD, n = 3).

*Process transfer to microfluidics*

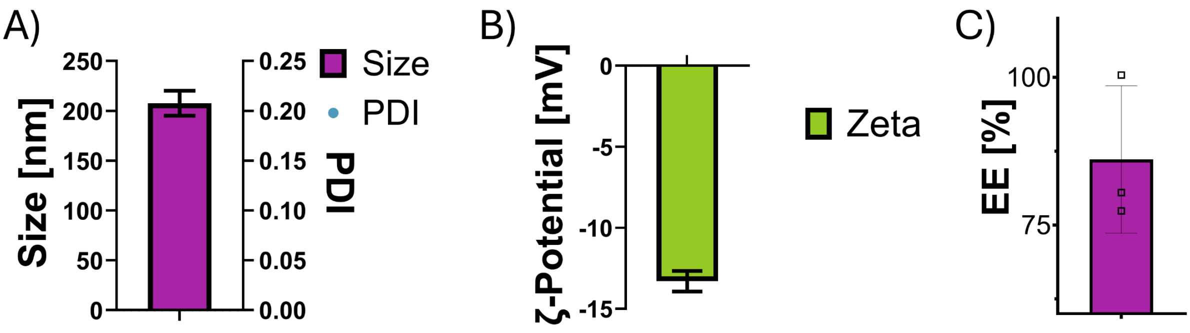

Fig S5 TILD System liposomes transferred to microfluidic production. A) Size and PDI, B) Zeta potential, and C) Encapsulation efficiency of TILD liposomes produced via microfluidics.

*In Vitro Drug Release*

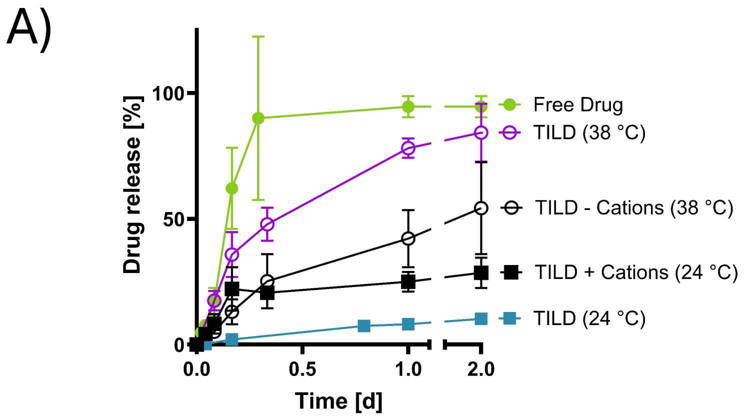

Fig S6 In vitro drug release of TILD system comparing free drug against TILD at different temperatures, and divalent cations to simulate the s.c. space and storage conditions.

*SAXS Spectra*

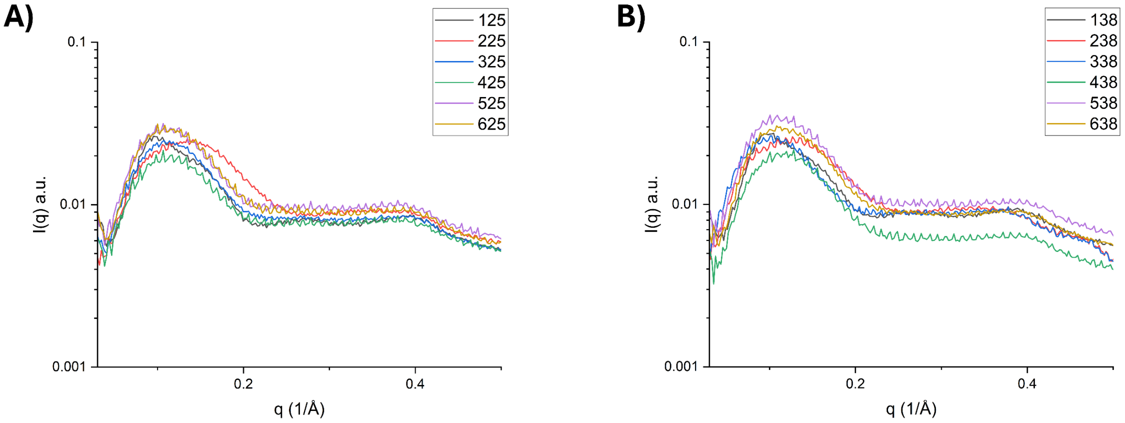

Fig S7 Scattering patterns of liposome measurements A) Measurements at 25 °C (125: HSPC; 225: DOPG; 325: HSPC/ DOPG (75:25 molar ratio) based-liposomes; 425: HSPC/ DOPG (75:25 molar ratio) based-liposomes + Buprenorphine (1:20); 525: HSPC/ DOPG (75:25 molar ratio) based-liposomes + 1 mM salt; 625: HSPC/ DOPG (75:25 molar ratio) based-liposomes + Buprenorphine (1:20) + 1 mM salt) B) Measurements at 38 °C (138: HSPC; 238: DOPG; 338: HSPC/ DOPG (75:25 molar ratio) based-liposomes; 438: HSPC/ DOPG (75:25 molar ratio) based-liposomes + Buprenorphine (1:20); 538: HSPC/ DOPG (75:25 molar ratio) based-liposomes + 1 mM salt; 638: HSPC/ DOPG (75:25 molar ratio) based-liposomes + Buprenorphine (1:20) + 1 mM salt)

*In vivo scoring*

Table S2 Scoring sheet during pharmacokinetic and pharmacodynamic study in rats.

| **Monitoring scale** | **Score standards** | **Details** |
|---|---|---|
| Demeanour | 0 | Active, interested |
| | 1 | Calm, moves when touched |
| | 2 | Immobile, apathic§ |
| Body weight | 0 | 0-9% weight loss from baseline |
| | 1 | 10-14% weight loss from baseline |
| Baseline BW = 186 | 2 | >15% weight loss from baseline§ |
| Appearance | 0 | Straight, bright pelt |
| | 1 | Dull, slightly ruffled fur |
| | 2 | Ruffled fur, hunched posture |
| Grimace scale | 0 | Open eyes and straight ears |
| | 1 | Orbital tightening, check bulge |
| | 2* | Orbital tightening, cheek bulge and flat ears§ |
| Paw appearance | 0 | Pink and soft |
| | 1 | Reddish and inflamed |
| | 2 | Red, inflamed and swollen |
| Paw inflammation | NA | Paw oedema (mm) |
| | 1 | Paw licking |
| | 1 | Paw guarding |
| Gastrointestinal | 1 | Diahrrea |
| | 2 | Pica§ |
| | 2 | Blood in faeces§ |
| Body T° (°C) | 2 | hypothermia§ |
| Baseline T°= | 2 | Hyperthermia§ |
| **Total Score** | Best= 0, Worst=16. | |

Absolute Cut offs§: Weight-loss of > 15%, blood in faeces, pica, hypothermia (T°<34°C), severe pain*, immobile and apathic

Table S3 Scoring sheet during pharmacokinetic study in dogs.

| **Monitor Scale** | **Score** | **Description** |
|---|---|---|
| Overall appearance | 0 | No sedation apparent |
| | 1 | Mild sedation |
| | 2 | Moderate sedation |
| | 3 | Well sedated |
| Posture | 0 | Standing |
| | 1 | Sitting |
| | 2 | Sternal recumbency |
| | 3 | Lateral recumbency |
| Awareness | 0 | Conscious and spontaneous interest |
| | 1 | Conscious but disinterested |
| | 2 | Unconscious |
| Interactive behaviour | 0 | Responds to 1 vocal stimulus |
| | 1 | Responds to 1 touch |
| | 2 | Does not respond to 1 touch |
| Spontaneous behaviour | 0 | Moves spontaneously |
| | 1 | Relaxed no tremors |
| | 2 | Very relaxed |
| Eye position | 0 | No rotation (normal) |
| | 1 | Moderate rotation |
| | 2 | Total rotation |
| Palpebral reflex<br>Awareness | 0 | Brisk |
| | 1 | Slow |
| | 0 | Conscious and spontaneous interest |
| **Total Score** | Lowest= 0, Highest= 25 | |

*Observation times: 1st Day: 1, 4 and 8 h after injection; consecutive days once per day from 8-9 am.

**Report score and exact measured value
